# Polynucleotide Phosphorylase Stops Replication Restart by Degrading the RNA within R-loops during Replication-Transcription Conflicts

**DOI:** 10.64898/2026.06.02.729377

**Authors:** Oyku Sensoy, Juan Carvajal-Garcia, Shahin Boumi, Houra Merrikh

**Author notes:** Co-first author (Oyku Sensoy, Juan Carvajal-Garcia). **Author Contributions:** O.S., J.C.-G. and H.M. designed research; O.S., J.C.-G., H.M., and S.B. performed research; O.S., J.C.-G., and H.M. analyzed data; and J.C.-G. and H.M. wrote the paper. **Competing Interest Statement:** None declared.

## Abstract

DNA replication and transcription machineries function simultaneously, on the same template, leading to conflicts between the two. During conflicts, the nascent mRNAs can hybridize to their template strands, generating RNA:DNA hybrids, which reside within nucleic acid structures known as R-loops. Accumulation of R-loops can lead to severe replication fork stalling, eventually leading to cell death. In bacteria, there are well-known replication restart mechanisms that should rescue these stalled replication forks, and yet, during conflicts, this process is inhibited. Additionally, in vitro work has shown that RNA:DNA hybrids are substrates for replication restart. Here, we discovered that the highly conserved exonuclease Polynucleotide Phosphorylase (PNPase in *Bacillus subtilis*) prevents replication restart from RNA:DNA hybrids, specifically at regions of severe conflicts where R-loops accumulate. Our in vitro data show that PNPase binds RNA:DNA hybrids and digests the RNA in these structures. Consistently, we found that in vivo, PNPase binds to conflict regions and reduces R-loop levels. To our knowledge, the only other class of enzymes known to digest the RNA from hybrids are RNases H. Our findings identify PNPase as a new enzyme that can perform a similar function. Importantly, our data show that PNPase inhibits replication restart from R-loops and its absence allows for replication restart from conflict regions. We also observed that PNPase activity reduces mutations, suggesting that replication restart from RNA:DNA hybrids is highly mutagenic. We propose that PNPase acts as a safeguard against mutagenic replication restart from RNA:DNA hybrids within R-loops by digesting the RNA which could re-initiate replication.

## Introduction

DNA replication is initiated from an RNA primer annealed to DNA (1). During replication elongation, such primers are used for synthesis of Okazaki fragments (2, 3). However, RNA:DNA hybrids can form outside of replication forks during various stages of transcription (4, 5). In these cases, the newly synthesized RNA can anneal to its template strand, displacing the non-template strand, and forming three-stranded nucleic acid structures known as R-loops (6). Although not always toxic (7), the RNA:DNA hybrids within the R-loop structures must be eliminated from the genome to ensure that DNA replication does not initiate inappropriately outside of canonical origins (8, 9). This can generally be achieved by the degradation of the RNA strand within R-loops.

R-loops are structures that can form in all species, from bacteria to humans, and their negative consequences are equally conserved (10, 11). They are a threat to genome stability (11), as they slow down or stall replication (12–14), promote mutagenesis (15, 16), and lead to DNA breaks (17, 18). For this reason, all organisms have enzymes that can resolve R-loops, which are essential to avoid the accumulation of these potentially pathological structures (18, 19).

The known and best understood R-loop resolution factors are the RNase H family of enzymes. These are RNA endonucleases that degrade RNAs that are annealed to DNA, including within R-loops (20). In bacteria, two enzymes in this family perform the bulk of R-loop resolution, RNase HI and RNase HIII, which are often mutually exclusive (21). In their absence, cells accumulate R-loops and the issues associated with these structures are exacerbated (19, 22). However, it remains unclear whether there are other enzymes outside of the RNase H family that can degrade the RNA strand within R-loop structures.

Polynucleotide phosphorylase (PNPase) is a highly conserved 3’ exonuclease that catalyzes the phosphorolysis of nucleic acids (23), especially RNA (24), but under certain conditions also DNA (25, 26). Additionally, PNPase catalyzes the reverse reaction: template-independent synthesis of RNA from ribonucleotide diphosphates (23), although its nuclease activity is considered to be the predominant function in vivo (27). PNPase is highly conserved and plays a key role in RNA metabolism in both bacteria and eukaryotes (27). Moreover, PNPase has been shown to play a role in mutagenesis and DNA repair in bacteria (26, 28, 29). The underlying mechanism of PNPase that leads to these consequences, especially mutagenesis, remains elusive.

Given its ability to degrade RNA and DNA, we wondered if PNPase functions at R-loop structures. We tested our hypothesis in *Bacillus subtilis* and found that PNPase can bind RNA:DNA hybrids in vitro. Furthermore, we found that in vivo, PNPase associates with regions where R-loops accumulate, specifically, at regions where the replication machinery encounters RNA Polymerases, leading to conflicts between the two. We determined that PNPase can degrade RNA:DNA hybrids at replication-transcription conflict regions, and that it specifically degrades the RNA strand of these structures.

Strikingly, the ability of PNPase to degrade RNA:DNA hybrids in vivo does not promote cell survival when cells experience severe head-on conflicts, which promote R-loop formation (18, 19), but rather leads to cell death, as cells lacking both RNase HIII and PNPase are able to survive R-loop accumulation significantly better than cells lacking only RNase HIII. Furthermore, cells lacking both enzymes have a much higher mutation frequency, and this is due to R-loop formation. This suggests that, while allowing cells to survive R-loop dependent replication stalling, the restart pathway that depends on the absence of the PNPase pathway is error-prone, which explains why cells have multiple pathways to suppress it. We propose that this stems from PNPase degrading RNA primers that can be used to re-start replication after it stalls due to R-loop accumulation (30). Taken together, these results highlight a non-canonical role for PNPase in R-loop degradation, suppressing error-prone replication restart, which comes at the cost of cell death when R-loops accumulate.

## Results

### Bacillus subtilis PNPase can bind and degrade RNA:DNA hybrids

Due to the ability of bacterial polynucleotide phosphorylase (PNPase) to bind and degrade both ssRNA and ssDNA (23–26, 31), we wondered if it is able to bind RNA:DNA hybrids. We expressed and purified *B. subtilis* PNPase (SI Fig. 1) and tested whether it can bind RNA:DNA hybrids in vitro by electrophoretic mobility shift assays (EMSAs) in vitro. We incubated PNPase with Cy5 labeled 47 nt ssRNA or ssDNA, or a 47 base pair hybrid, with the same sequences, and ran them on a non-denaturing polyacrylamide gel. We observed a clear shift in the position of all three types of nucleic acids we constructed (Fig. 1a), indicating stable binding of PNPase to the substrates. These results clearly show that PNPase binds RNA:DNA hybrids.

**Figure 1.**
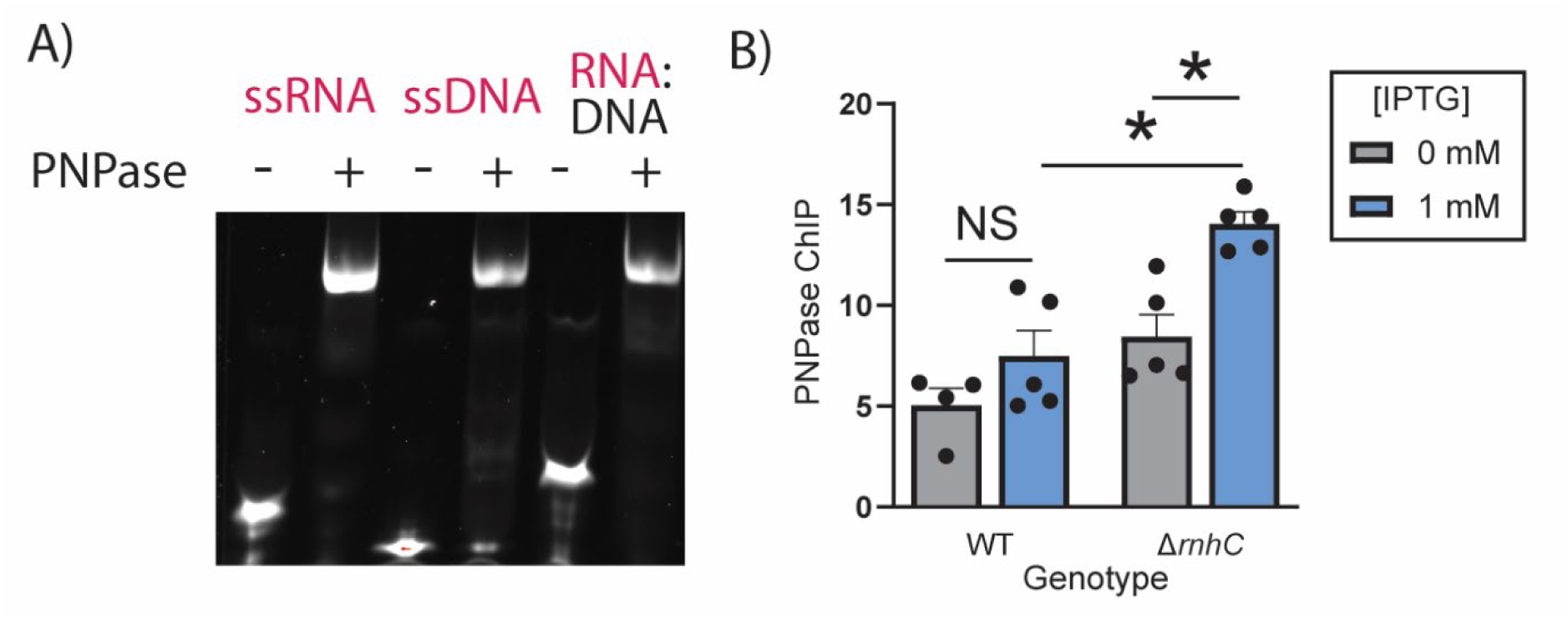
PNPase binds RNA:DNA hybrids. A) EMSA of WT PNPase and the indicated substrates. The labeled strand in the substrate is indicated in red. B) ChIP-qPCR of myc-tag PNPase in the indicated strain indicated as % IP. Error bars represent the standard error of the mean. Statistical significance was assessed by one-way ANOVA. \**p<*0.05.

To test whether PNPase binds regions where there is R-loop accumulation in vivo, we added three myc-tags to the 3’ end of the endogenous *pnpA* gene. Using an engineered head-on replication-transcription conflict system in cells, which we have shown accumulates RNA:DNA hybrids in the form of R-loops (30), we explored whether PNPase binds to these regions by chromatin immunoprecipitations (ChIPs).The engineered conflict consists of the *hisC* gene integrated in the head-on orientation to replication, into the *amyE* locus, under the control of the strong inducible promoter *Pspank*(*hy*). Addition of IPTG to the media turns on the expression of *hisC*, leading to severe replication-transcription conflicts (19). While we did not observe transcription-dependent PNPase recruitment to the conflict region in WT cells, we did observe it in cells that were lacking the well-known R-loop resolution protein in *B. subtilis*, RNase HIII (Fig. 1b). This result suggested that PNPase might serve as a secondary R-loop resolution factor.

To test this, we determined the RNA:DNA hybrid levels at the conflict region by DNA-RNA immunoprecipitation (DRIP) followed by qPCR in WT cells as well as Δ*pnpA*, Δ*rnhC*, and Δ*pnpA* Δ*rnhC* double mutants. We have shown in the past that Δ*rnhC* cells show an accumulation of R-loops at sites of replication-transcription conflicts (19, 30). Strikingly, we observe a similar R-loop accumulation in Δ*pnpA* cells, and an additive effect of the double mutant Δ*rnhC* Δ*pnpA* (Fig. 2a-b). Interestingly, by looking at R-loop levels over time, we observed that Δ*rnhC* cells rapidly accumulate R-loops, but this is followed by a decrease in their R-loop levels, suggesting that PNPase is responsible for degradation of these structures in the absence of RNase HIII, albeit with slower kinetics (SI Fig. 2). Consistent with this, we do not see this time-dependent decrease in Δ*pnpA* Δ*rnhC* double mutant cells (SI Fig. 2). Rather, we observed continued R-loop accumulation over time, which led us to conclude that PNPase decreases R-loops levels in vivo.

**Figure 2.**
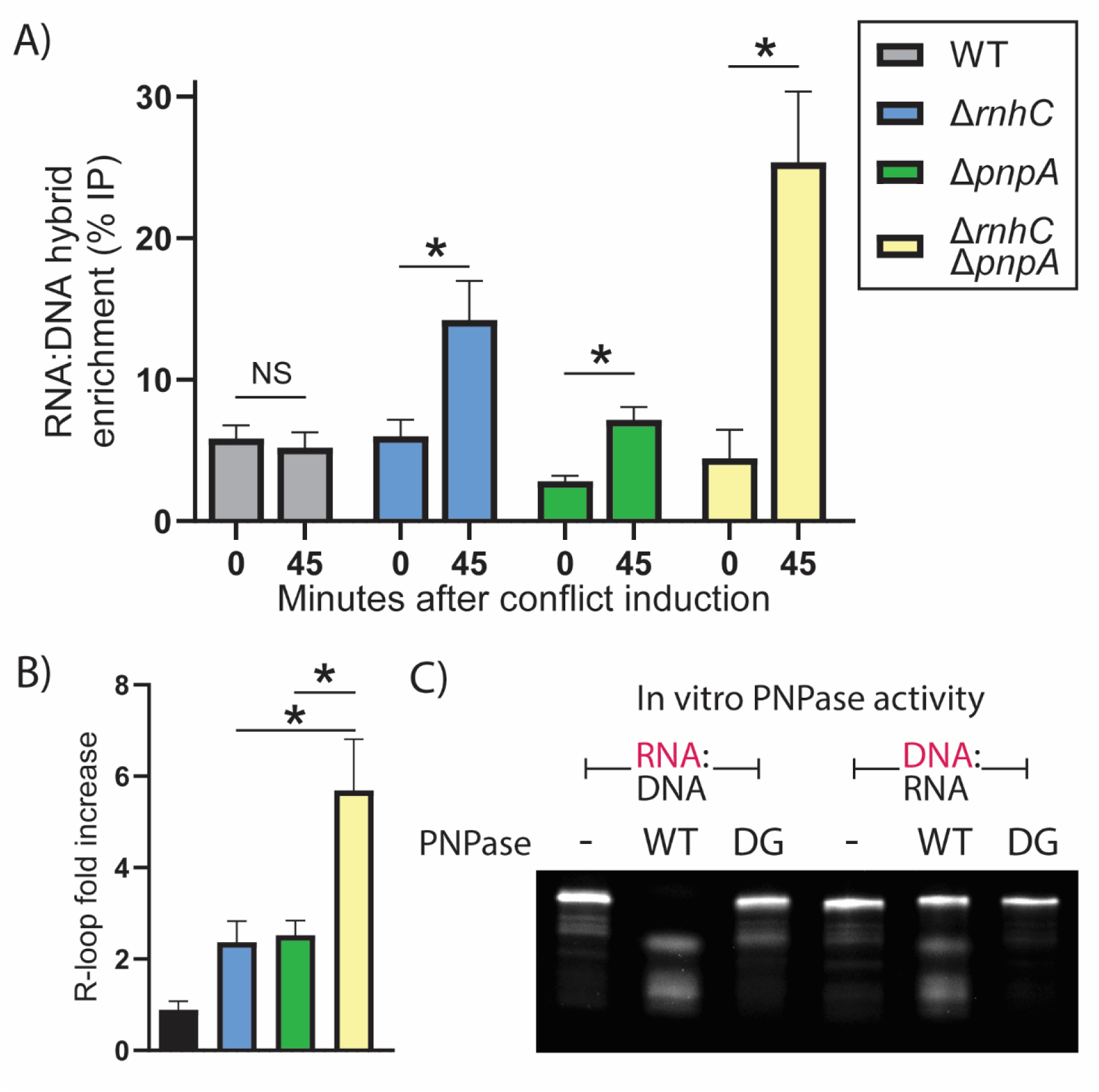
PNPase degrades the RNA strand of RNA:DNA hybrids. A) DRIP in the indicated strain represented as % IP before and 45 minutes after conflict induction. Error bars represent the standard error of the mean. Statistical significance was assessed by two-tailed t-test. \**p<*0.05. B) Fold increase in DRIP signal 45 minutes after conflict induction, calculated using the data from panel A. Error bars represent the standard error of the mean. Statistical significance was assessed by two-tailed t-test. \**p<*0.05. C) Representative 10% urea gel of the indicated substrates after incubation at 37 °C for 5 minutes with either WT or D493G (DG) PNPase in the presence of 10 µM phosphate. The labeled strand in the substrate is indicated in red.

We set out to determine if the observed R-loop accumulation in the absence of PNPase might be due to this protein being able to degrade RNA:DNA hybrids. To do so, we tested the ability of PNPase to degrade an RNA:DNA hybrid in vitro by incubating the protein with an RNA:DNA hybrid in the presence of phosphate, which is required for PNPase’s exonuclease activity. For this, we used two hybrids, one with the RNA strand labeled, and one with the DNA strand labeled, and observed that PNPase degrades the RNA strand of an RNA:DNA hybrid much more efficiently than the DNA strand (Fig. 2c). Importantly, we did not observe any RNA or DNA strand degradation in a mutant PNPase (D493G) that has been shown in the past to be catalytically dead (Fig. 2c) (29).

### PNPase prevents cell survival in the absence of RNase HIII

The additive effect of RNase HIII and PNPase in R-loop degradation led us to hypothesize that they might have a similar additive effect in the survival of cells experiencing severe head-on conflicts. To test this, we used the engineered *hisC* conflict system, where strong expression of this gene by IPTG leads to cell death in cells lacking RNase HIII. However, lower concentrations of IPTG, which lead to reduced expression of the inducible *hisC* gene, allow us to observe a partial survival of these cells and therefore to test our hypothesis using genetics.

We measured the survival of WT, Δ*pnpA*, Δ*rnhC*, and Δ*pnpA* Δ*rnhC* cells under various levels of conflict induction using increasing concentrations of IPTG. Strikingly, we observed that contrary to our expectation, lack of PNPase almost completely rescues the death of Δ*rnhC* cells, even at the highest level of conflict induction (Fig. 3a).

**Figure 3.**
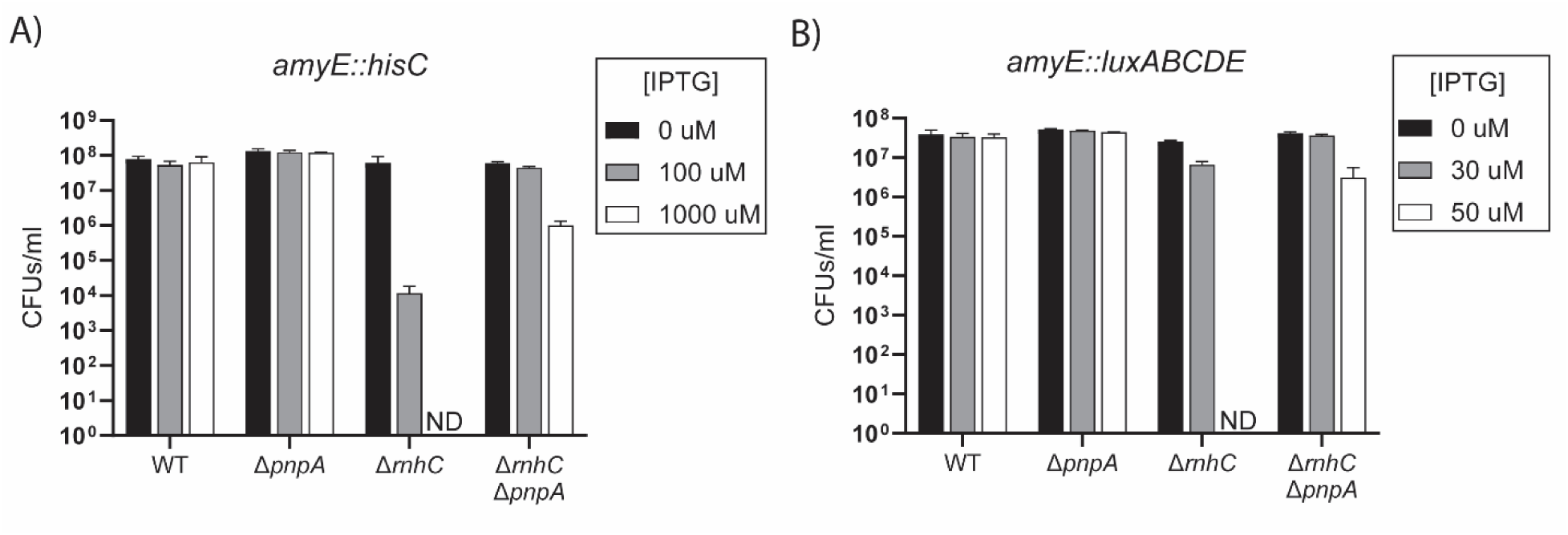
PNPase leads to cell conflict-dependent cell death. A-B) Colony forming units/ml at an OD600 of 0.3 of the indicated strains plated in the presence of the indicated concentration of IPTG. ND=not detected.

To make sure that this effect is not exclusive to the *hisC* gene, we used a different engineered conflict in which the *luxABCDE* operon is introduced into the *amyE* locus, under the control of the same *Pspank(hy)* promoter. We observed a similar rescue using this conflict, although we had to use a lower concentration of IPTG (Fig. 3b). This is likely due to the fact that replication-transition conflicts caused by longer genes are more severe than those caused by shorter ones. *hisC* is 1 kb in length whereas the *luxABCDE* operon is approximately 5.5 kb – by definition, leading to replication-transcription conflicts more often.

The rescue of viability in Δ*pnpA* Δ*rnhC* cells with a severe head-on conflict suggests that excessive degradation of R-loops by PNPase might be responsible for the cell death observed in Δ*rnhC* cells. However, an alternative hypothesis is that the lack of PNPase might be decreasing the severity of the conflict, allowing replication to proceed seamlessly past the conflict region. To test this, we determined the genome copy number of these cells by high throughput sequencing during exponential growth (Fig. 4). In cells experiencing a severe head-on conflict, and lacking conflict resolution factors, this type of analysis reveals that replication cannot proceed past the conflict region (19, 32), which is observed as an abrupt decrease in the number of sequencing reads that align to the regions downstream of the conflict region and up until the terminus. If the lack of PNPase was leading to a less severe conflict, we would not see such a decrease in Δ*pnpA* Δ*rnhC* cells. However, we observe that replication stalls just as significantly as we have previously observed in Δ*rnhC* cells (19) in the double mutants after the conflict is induced, while no stalling is observed in WT or Δ*pnpA* cells (Fig. 4).

**Figure 4.**
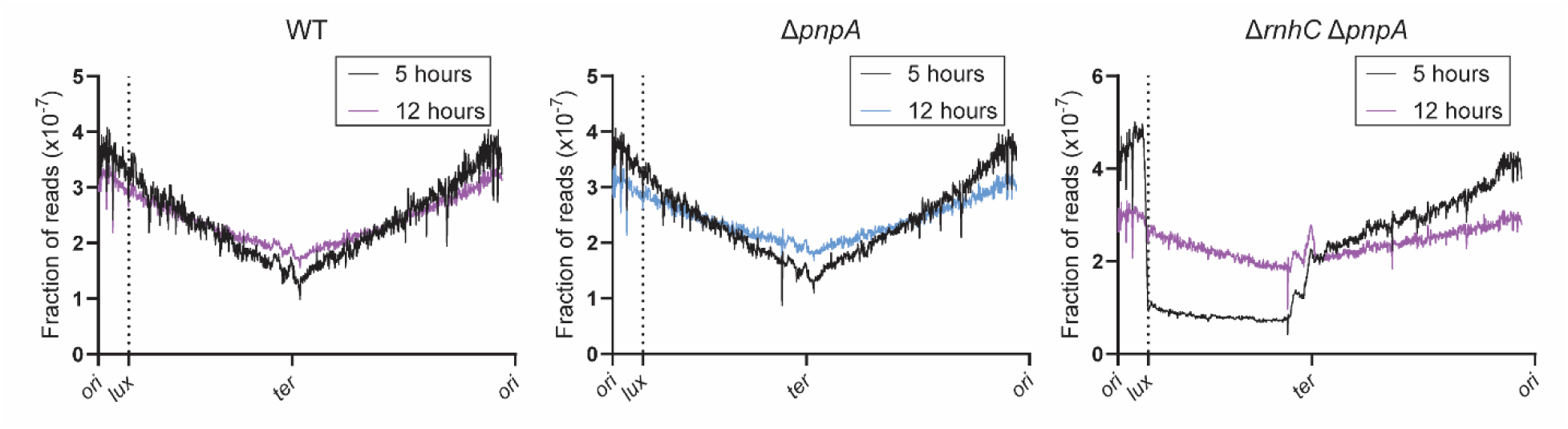
PNPase prevents replication restart from a conflict region. Fraction of sequencing reads that map to the indicated coordinates (5 kb window) of the *B. subtilis* genome, smoothed using a second other polynomial using 10 neighbors on each side in the indicted strains grown at 37 °C for the indicated time after conflict induction by addition of 1 mM IPTG.

Importantly, these cells are viable, which means that the sequences downstream of the conflict have to be replicated eventually. Accordingly, when we grow the cells in liquid media and check their optical density over time, we see that the growth of Δ*pnpA* Δ*rnhC* double mutants are delayed when compared to WT or Δ*pnpA* cells, but they eventually start growing (SI Fig. 3a). When we determined the genome copy number in Δ*pnpA* Δ*rnhC* cells 12 hours after the induction of the conflict, we see that the region downstream of the conflict is being replicated, as there are more sequencing reads in this region than in the earlier timepoint (Fig. 4). Our interpretation of this data is that Δ*pnpA* Δ*rnhC* cells, albeit slowly, are able to restart replication from the conflict region, allowing cell survival despite the initial transcription-induced stalling of the replication fork.

The experiments described above were performed in cells where we have engineered conflicts, but these conflicts can naturally occur in native genes, especially in those that are expressed head-on to replication. An example of this is the *dltABCDE* operon, which is highly induced by lysozyme (19, 33), and is in head-on orientation with respect to replication. When *B. subtilis* grows in the presence of subinhibitory concentrations of lysozyme, this leads to replication-transcription conflicts that require the R-loop resolution factor RNase HIII for survival (19). Consistent with our conclusions from the data described above, where PNPase is predicted to stall replication restart from head-on conflict regions, we observed that cells lacking PNPase are able to survive higher amounts of lysozyme than WT cells (SI Fig. 3b). This suggests the role of PNPase in preventing replication restart from head-on conflict regions is relevant in nature and not restricted to artificially engineered conflicts. Furthermore, these results show that even in the presence of RNase HIII, PNPase plays a role in regulating replication restart, establishing the importance of its function in R-loop resolution.

### Replication restart from an RNA:DNA hybrid is mutagenic

R-loops are known to be mutagenic (18, 19). To determine if PNPase function in R-loop resolution impacts mutagenesis, we measured mutation frequencies in WT, Δ*pnpA* and Δ*rnhC* Δ*pnpA* cells using the engineered *hisC* conflict. This *hisC* allele, *hisC952*, has a mutation that leads to a premature stop codon. A mutation that changes this stop codon to a coding one makes for a functional gene. Deleting the endogenous *hisC* gene allowed us to find these mutations and determine mutation frequencies, as only the cells harboring a reversion of the stop codon in *hisC952* can grow in minimal media lacking histidine. We cannot measure mutagenesis in Δ*rnhC* cells alone as they are not viable in the face of head-on conflicts such as during the expression of the *hisC* gene in these experiments.

We saw a large increase in the mutation frequencies in the Δ*rnhC* Δ*pnpA* double mutants compared to either WT or Δ*pnpA* single mutant cells (Fig. 5, SI Table 1). Importantly, overexpressing WT RNase HIII, which we have shown in the past decreases the levels of R-loops in cells (19), decreases mutations in all three genotypes (Fig. 5, SI Table 1). This shows that R-loops are responsible for the increase in mutation frequencies caused by the absence of PNPase. These data show that, while removing PNPase allows cells to survive, it does so by allowing a highly mutagenic pathway to take over, which we postulate is the type of replication restart discussed here. In other words, these data, together with the rest of the described observations presented here, suggest that PNPase normally suppresses a highly mutagenic replication restart pathway.

**Figure 5.**
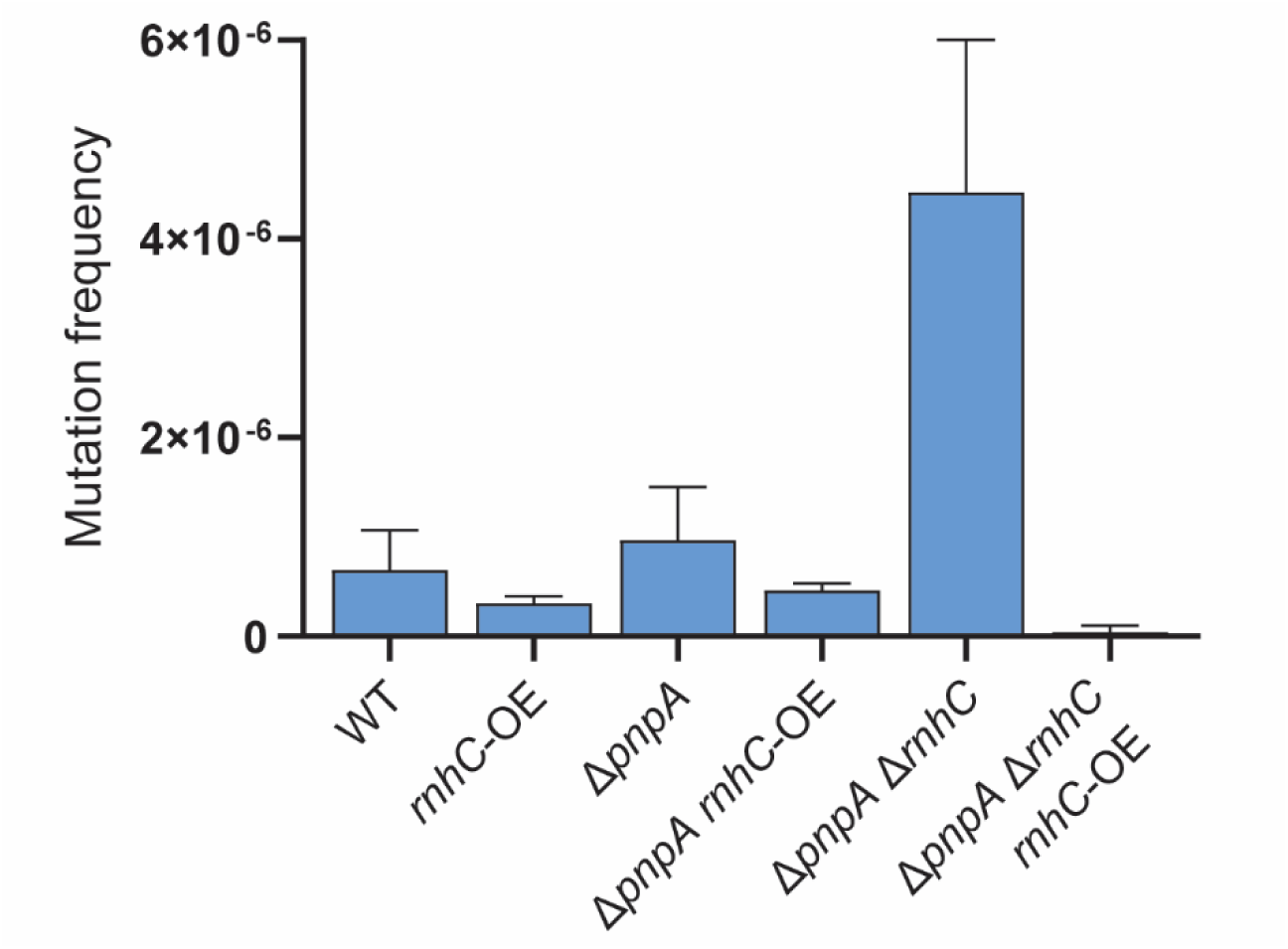
PNPase promotes R-loop dependent mutagenesis. Median mutation frequencies of the indicated strains determined using *hisC952* reversion. Error bars represent 95% confidence intervals. N=≥38 biological replicates.

## Discussion

DNA polymerases involved in replication require an RNA primer annealed to DNA to start synthesis (1). However, transcription also leads to RNA:DNA hybrids when the mRNA anneals to its template DNA strand, forming R-loops (7). R-loops can be formed when replication and transcription encounter each other at a replication-transcription conflict site (9, 18, 19), and need to be removed for further progression of the replication fork. R-loops can also form during RNA Polymerase backtracking or termination.

Up until this work, one family of enzymes had been described that is able to specifically degrade the RNA strand on an RNA:DNA hybrid, the RNase H family of endonucleases (20). Here we found that PNPase is also able to specifically remove the RNA strand of a hybrid, in this case by exonucleolytic phosphorolysis. This activity translates to an ability to resolve R-loops in general and especially those that are generated due to replication-transcription conflicts in vivo. Therefore, this study identifies PNPase as the second type of enzyme that can bind and resolve RNA:DNA hybrids by digesting the RNA strand. Given the high level of conservation found in nature, this work suggests that PNPase is a universal R-loop resolution factor.

However, removal of RNA:DNA hybrids by PNPase does not lead to an improvement in survival in cells that are experiencing severe head-on conflicts, but rather an increase in cell death. We observed that cells lacking the main R-loop resolution protein RNase HIII, which die when R-loops accumulate, are able to survive in the absence of PNPase. Furthermore, the removal of PNPase promotes survival in cells that are growing in the presence of lysozyme, which leads to replication-transcription conflicts. We hypothesize that the reason behind these paradoxical results is that PNPase, by degrading the RNA strand, is preventing replication restart from the R-loop which is likely deleterious to cells (Fig. 6). We predict that this adverse effect is due to the high mutational burden caused by restart events utilizing RNA:DNA hybrids. In essence, cells rather sacrifice themselves that be burdened by increased mutagenesis caused by RNA:DNA hybrid primed replication restart.

**Figure 6.**
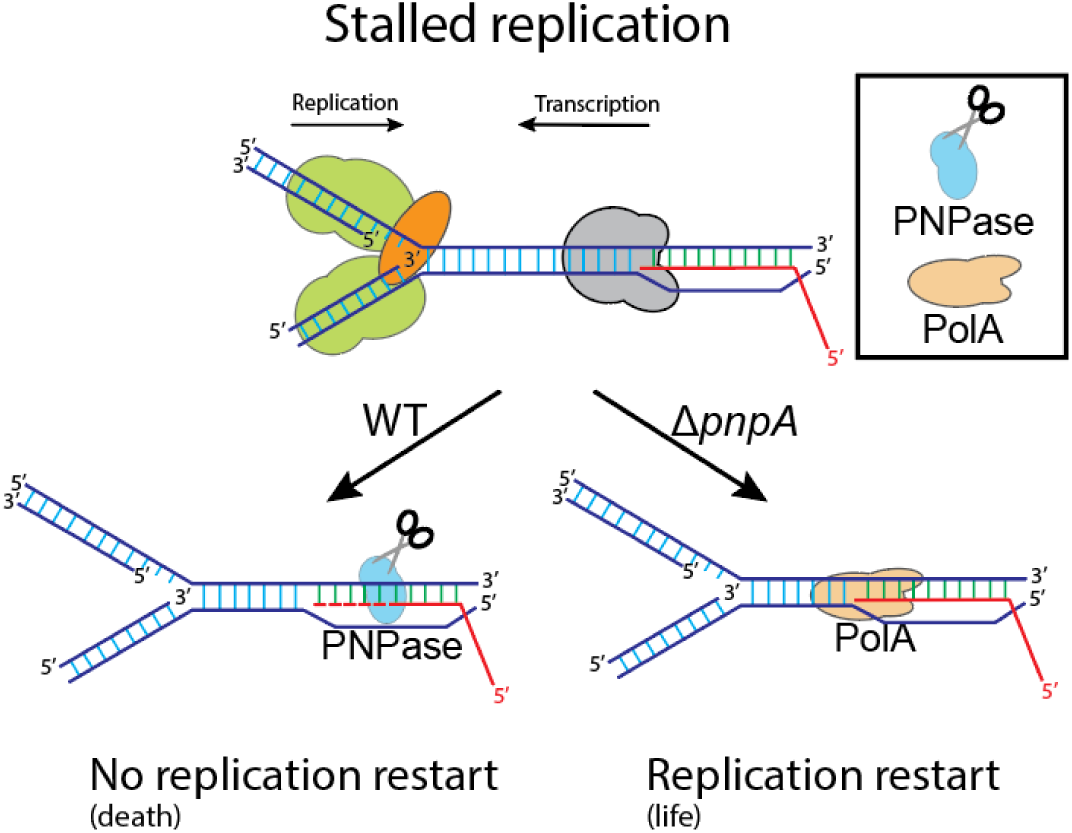
Model for PNPase-suppressed replication restart. After replication stalls due to a highly transcriptive head-on gene, the RNA from the RNA:DNA hybrid is degraded by PNPase. In the absence of PNPase, PolA (or another error prone DNA Polymerase) can use this hybrid to continue synthesis, allowing the cells to survive while increasing mutation levels.

Replication restart from the RNA strand of an RNA:DNA hybrid has been observed both in vitro and in vivo (8, 9, 34). Our recent findings in another study shows that restart from hybrids is highly regulated (30). In our previous work, we observed that hybrid-dependent replication restart depends on DNA polymerase I (PolA) (Fig. 6) and the DNA translocase Mfd, which removes RNA Polymerases from DNA, exposing the 3’ end necessary for DNA synthesis. We found that replication restart is blocked at conflict regions, where R-loops have accumulated, through the capping of the 3’ end of RNA by the asparagine tRNA ligase AsnRS (30). Here, we unravel a second regulatory mechanism for restarting replication from RNA:DNA hybrids: degradation of the RNAs present in R-loops by PNPase (Fig. 6).

By generating whole-genome replication profiles, we are able to infer that the mechanism we are observing is due to the inhibition of replication restart. Replication first stalls completely at the conflict region, showing that in the absence of PNPase conflicts still occur. But post this event, following the conflict, Δ*rnhC* Δ*pnpA* double mutant cells can restart replication, although this process is relatively slow.

We have found that there are at least two ways to prevent replication restart from an R-loop. This level of negative regulation suggests that, under normal conditions, restarting replication from RNA:DNA hybrids outside of Okazaki fragments is disfavored by cells. We recently showed that the most commonly used pathway for continuing replication after head-on replication conflicts is likely fork reversal and remodeling by the helicase-nuclease complex AddAB, which is not mutagenic (32, 35).Therefore, given all of our studies together, we suggest that restart upon head-on conflicts through fork remodeling is highly beneficial as it avoids restart initiated from R-loops, avoiding fork-fork collisions, general genome instability, and excessive mutagenesis, all of which would be highly deleterious to cells.

## Materials and Methods

### Bacterial culture

*Bacillus subtilis* was cultured in lysogeny broth (LB). In plates, cells were grown at 37 °C with the following antibiotics when appropriate: 500 µg/ml erythromycin and 12.5 µg/ml lincomycin (MLS),100 µg/ml spectinomycin, 5 µg/ml (*B. subtilis*) or 50 µg/ml (*E. coli*) kanamycin, and 100 µg/ml carbenicillin. Single colonies were used to start liquid cultures and were grown with aeration (260 rpm). A list of all strains used in this study can be found in SI Table 2 and a list of plasmids can be found in SI Table 3.

### Strain and plasmid construction

HM1 (also known as AG174 or JH642) (36, 37) is the parental strain for all *B. subtilis* strains used. MLS and kanamycin resistance-marked single-gene deletions were obtained from (38), and transformed into HM1 using standard protocols (39). When indicated, the antibiotic-resistant markers were excised by transforming the strains with a plasmid expressing the Cre recombinase (pDR244, BGSCID: ECE274) purified from Rec+ *E. coli* cells (AG1111) (40) generating markerless strains.

For PNPase expression and purification the coding sequence of *pnpA* minus the first nucleotide was amplified (SI Table 3) and cloned into the BamHI site of pET28a (Thermo) using NEBuilder® HiFi DNA Assembly Master Mix (NEB) generating plasmid pHM786 (SI Table 3). To generate the D493G mutant protein, pHM786 was amplified with 5’ phosphorylated primers, one of which contained the appropriate mutation (SI Table 3), and the PCR product was circularized with T4 DNA ligase (NEB), creating pHM787.

For adding 3 myc tags to PNPase, approximately 600 bp of the 3’ of the gene were amplified (SI Table 3) and cloned into pCAL812 (41). The spectinomycin resistance gene in these plasmids was changed to an MLS resistance amplified from *mfd*::MLS (HM2521) (35) using NEBuilder® HiFi DNA Assembly Master Mix (SI Table 2) and generating pHM795. This plasmid, purified from AG1111 cells, was transformed into competent *B. subtilis* cells.

### PNPase purification

WT and D493G PNPase were expressed and purified in parallel. pHM786 (WT) or pHM787 (D493G) were transformed into BL21(DE3)pLysS Competent Cells (Agilent), and a single colony was inoculated into 20 ml of LB and grown overnight in LB containing kanamycin. 10 ml of culture were used to inoculate 1 L of LB containing kanamycin, which was grown to an OD of 0.6, after which 1 mM IPTG was added to the media. The cultures were grown for an additional 4 hours, centrifuged for 15 mins at 6000G, and stored at -80 °C. The cells were resuspended in 10 ml of CelLytic B cell lysis reagent (Sigma) with 2 µl of Benzonase (Sigma), 10 mM imidazole and one cOmplete™, EDTA-free Protease Inhibitor Cocktail tablet (Millipore) and shaken gently at RT for 10 mins. The lysate was centrifuged at 20000G, 4 °C for 30 minutes and the supernatant was mixed with an equal volume of equilibration buffer (20 mM sodium phosphate pH 7.4, 300 mM sodium chloride, 10 mM imidazole), and run twice through 6 ml of equilibrated HisPur™ Ni-NTA Resin (Thermo) at 4 °C. The resin was washed with 50 ml of wash buffer (20 mM sodium phosphate pH 7.4, 300 mM sodium chloride, 40 mM imidazole) and eluted with 10 ml of elution buffer (20 mM sodium phosphate pH 7.4, 300 mM sodium chloride, 250 mM imidazole). The protein was dialyzed with a 15 ml Slide-A-Lyzer G3 Dialysis Cassette G2 20000 MWCO (Thermo) against 10 mM tris pH 8, 50 mM NaCl, 5% glycerol, 0.1 mM DTT, 0.1 mM EDTA overnight at 4 °C and concentrated with two Amicon Ultra-4 Centrifugal Filter Units 50000 NMWL (Millipore) to a concentration of 1.8 mg/ml (WT) and 2.4 mg/ml (D493G) measured by Qubit (Thermo). The protein preps were run in a 4-20% Mini-PROTEAN® TGX™ Precast Protein Gel (BioRad) and stained with Bio-Safe™ Coomassie Stain (BioRad) to confirm >95% purity (SI Fig. 1).

### Electrophoretic Mobility Shift Assays

1.5 nM of the indicated substrate (hybrid substrates were annealed on a thermocycler) was incubated in 1X rCutSmart buffer (NEB) with 400 nM PNPase when indicated for 5 minutes on ice. A 0.5X TBE, 6% native polyacrylamide gel made with was pre-ran for 30 minutes at 4 °C before adding glycerol to a final concentration of 6.66% and loading the samples, which were run at 4 °C for approximately 45 minutes. The gels were scanned in a ChemiDoc imaging system (BioRad).

### Chromatin Immunoprecipitation

A single colony of the indicated strain was inoculated into 5 ml of LB and grown to an OD600 of >0.5. This culture was then diluted to an OD600 of 0.05 in LB supplemented or not with the indicated amount of IPTG and grown to an OD of approximately 0.5. Cultures were then processed as in Merrikh et al., 2011. In short, cultures were crosslinked with 1% formaldehyde for 20 minutes at room temperature, and quenched with 0.5 M glycine for 5 minutes. The crosslinked cultures were centrifuged and washed with ice-cold 1X phosphate-buffered saline (PBS). The pellets resuspended in 1.5 ml of solution A supplemented with 1 mM AEBSF and incubated for 30 min at 37 °C, followed by lysis with 1.5 ml of 2X IP buffer (100 mM Tris pH 7.0, 10 mM EDTA, 2% triton X-100, 300 mM NaCl and 1mM AEBSF) and incubation on ice for 30 mins. The lysates were then sonicated 4 times at 30% amplitude alternating 10 s of sonication and 10 s of rest and centrifuged at 70000 G and 4 °C for 15 min. 1 ml of supernatant was incubated with 12 µg of 9E10 c-Myc Monoclonal Antibody (Thermo, MA1-980) on a mini-tube rotator at 10 rpm at 4 °C overnight (IPs). 40 µl of lysates to be used as inputs were diluted with 50 µl of TE pH=8 and SDS was added to a concentration of 1%. 30 µl of Protein G Sepharose™ 4 Fast Flow resin (Cytiva, 17061801) were washed with 1X IP buffer for 10 mins in a mini-tube rotator at 10 rpm at room temperature and added to the IPs, which were rotated for 1 h at 10 rpm at room temperature. The beads were washed 6 times with 1X IP buffer and once with TE pH=7.6, and eluted with 100 μl of elution buffer (50 mM Tris pH 8.0, 10 mM EDTA, 1% SDS) by incubating at 65 °C for 10 minutes. The beads were centrifuged and the supernatant was saved, followed by the addition of 150 μl of elution buffer II (10 mM Tris pH 8.0, 1 mM EDTA, 0.67% SDS) to the beads. The beads were again centrifuged, and the supernatant was combined with the first elution. The combined eluates were de-crosslinked by incubation at 65° C overnight. 0.4 mg/ml of proteinase K was then added to both the eluates and the inputs, which were incubated for 2 hours at 37 °C. The DNA from both the IPs and the inputs was purified by standard phenol:chloroform extraction followed by ethanol precipitation, and resuspended overnight with 75 µl of TE buffer pH=8. qPCR analysis was performed using SsoAdvanced Universal SYBR Green Supermix (Bio-Rad), in a CFX96 Touch Real-Time PCR system (Bio-Rad) thermocycler, the inputs were diluted 1:500 before adding to the qPCR reaction. Mean Ct values were determined separately for IP and input samples, and percent IP was calculated.

### DNA:RNA hybrid immunoprecipitation assays (DRIP)

DRIP assays were performed as previously described (42) with minor modifications (22). In brief, pre-cultures were grown from a single colony to the mid-exponential phase (OD600 of 0.4-0.7) at 37°C with shaking (260 rpm), diluted back to OD600 of 0.05 in LB medium (25 ml), and grown for the indicated time at 30°C in the presence of 1mM IPTG. Cells were centrifuged by centrifugation, washed in cold 1X PBS buffer, and resuspended in TE Buffer (pH 8) with 10 mg/ml lysozyme.

Cells were lysed at 37°C for 30 minutes and lysates were treated with Proteinase 20 mg/ml K at 37°C overnight. DNA was extracted from lysates by phenol-chloroform-isoamyl alcohol (25:24:1) and precipitated by ethanol precipitation. DNA was washed twice with 80% ethanol, and air dried at room temperature. DNA was resuspended in TE Buffer (pH 8) and then digested overnight at 37°C with HindIII, EcoRV, EcoRI, and DraI restriction enzymes (NEB). Digested chromosomal DNA was then extracted with phenol chloroform-isoamyl alcohol (25:24:1) followed by ethanol precipitation and resuspended with TE Buffer (pH 8). At this point, 10% of the DNA volume was collected as input control. The remaining DNA was incubated overnight at 4°C on a mini-tube rotator (10 rpm) in 10X DRIP Binding Buffer (100 mM NaPO4 pH 7, 1.4 M NaCl, 0.5% Triton X-100) with 20μl of the S9.6 antibody (Vanderbilt Antibody and Protein Resource (VAPR) Core). 30 µl of Protein A sepharose beads (GE) were added to IP samples and incubated at 4°C for 2 hours on a mini-tube rotator (10 rpm). Beads were washed twice in 1X DRIP Binding Buffer for 15 minutes at room temperature on a mini-tube rotator (10 rpm) and the DNA was eluted with 300 μl of DRIP Elution Buffer (10 mM Tris pH 8, 1mM EDTA, 0.67% SDS) followed by incubation at 55°C for 45 minutes. DNA was then extracted with phenol chloroform-isoamyl alcohol (25:24:1) followed by ethanol precipitation and resuspended with TE Buffer (pH 8). qPCR analysis was performed using SsoAdvanced Universal SYBR Green Supermix (Bio-Rad), in a CFX96 Touch Real-Time PCR system (Bio-Rad) thermocycler, the inputs were diluted 1:500 before adding to the qPCR reaction. Mean Ct values were determined separately for IP and input samples, and percent IP was calculated.

### In vitro PNPase degradation assays

1.5 nM of the indicated substrates were mixed in 1X rCutSmart buffer and 10 µM ammonium phosphate on ice. The reactions were started by adding the indicated protein at a concentration of 10 nM, and were incubated at 37 °C for 5 minutes, after which they were stopped with by adding an equal volume of 98% formamide 10 mM EDTA. The substrates were denatured at 98 °C for 5 min and ran in a 10% Mini-PROTEAN® TBE-Urea Gel (BioRad). The gels were scanned in a ChemiDoc imaging system (BioRad).

### Survival assays

A single colony of the indicated strain was grown on 2 ml of LB to the mid-exponential phase (OD600 of 0.4-0.7) at 37°C with shaking (260 rpm). These cultures were diluted back to OD600 of 0.3 and serially diluted (1:10) in 1X Spizizen’s Minimal Salts solution. 5μl of each dilution was plated onto LB agar plates with the indicated concentrations of IPTG. Plates were incubated overnight at 30°C and colonies were counted.

### Lysozyme MIC determination

A single colony of the indicated strain was grown in 2 ml of LB to mid-exponential phase (OD600 of 0.4-0.7) at 37°C with shaking (260 rpm). Pre-cultures were diluted back to OD600 of 0.0025 in a total volume of 200μl using fresh LB in 96 well plates with the indicated concentration of freshly dissolved lysozyme. The plates were grown at 37°C with shaking (260 rpm) for 16 hours and the OD was determined with a Epoch2 plate reader. Each biological replicate represents the average of 8 technical replicates.

### Growth curve determination in liquid media

A single colony of the indicated strain was grown in 2 ml of LB to mid-exponential phase (OD600 of 0.4-0.7) at 37°C with shaking (260 rpm). Cells were diluted to OD600 of 0.0025 in a total volume of 200μl of LB with 1 mM IPTG in clear 96-well plates. Plates were then incubated in an Epoch2 plate reader at 37°C with shaking, and the OD was determined every 10 minutes.

### Replication profiling

Cells were prepared and grown as in “Growth curve determination in liquid media”. Genomic DNA from the indicated strains 5 and 12 hours of after conflict induction was purified with the GeneJET Genomic DNA Purification Kit (thermo). The genomic DNA was submitted to the Vanderbilt Technologies for Advanced Genomics (VANTAGE) where it was used to prepare Illumina libraries that were sequenced in a NovaSeq XP Plus (2×150 flow cell). Low quality treads and bases were trimmed with fastp (43), and the resulting reads were aligned to the appropriate strain using bowtie2 (44).

### Fluctuation tests

A single colony from the indicated strain was grown for 14 hours in 2 ml of LB with spectinomycin. Then, the culture was diluted into 12 parallel cultures to an OD600 of 0.0005 in 2 ml of LB with 1 mM IPTG and grown to an OD600 of 0.5-0.7. 100 µl of culture were serially diluted and plated on Spizizen’s Minimal Medium (0.2 mg/ml ammonium sulphate, 1.4 mg/ml monobasic potassium phosphate, 0.6 mg/ml dibasic potassium phosphate, 0.1 mg/ml sodium citrate dihydrate, 0.02 mg/ml magnesium sulphate heptahydrate, 100 μg/ml glutamic acid, 5 μg/ml glucose) supplemented with 40 µg/ml of phenylalanine, tryptophane, threonine, methionine, leucine and histidine and 1 mM IPTG to determine the number of viable cells. 1.5 ml of culture were centrifuged, re-suspended in Spizizen’s Minimal Salts, and plated on Spizizen’s Minimal Medium supplemented with the amino acids below expect for histidine and 1 mM IPTG to detect revertant colonies. All plates were grown at 37 °C and colonies were counted after 36-48 hours.

## Acknowledgements

This work was supported by K99ES037493 to JC-G and Vanderbilt University School of Medicine Department of Biochemistry funds to HM.

This research was also supported by the Vanderbilt School of Medicine Basic Sciences, Department of Biochemistry, and the Destination Biochemistry Advanced Postdoctoral Scholar Award funded by the Armstrong Family to JC-G.

## Supplementary Information

### SI Figures

**SI Figure 1.**
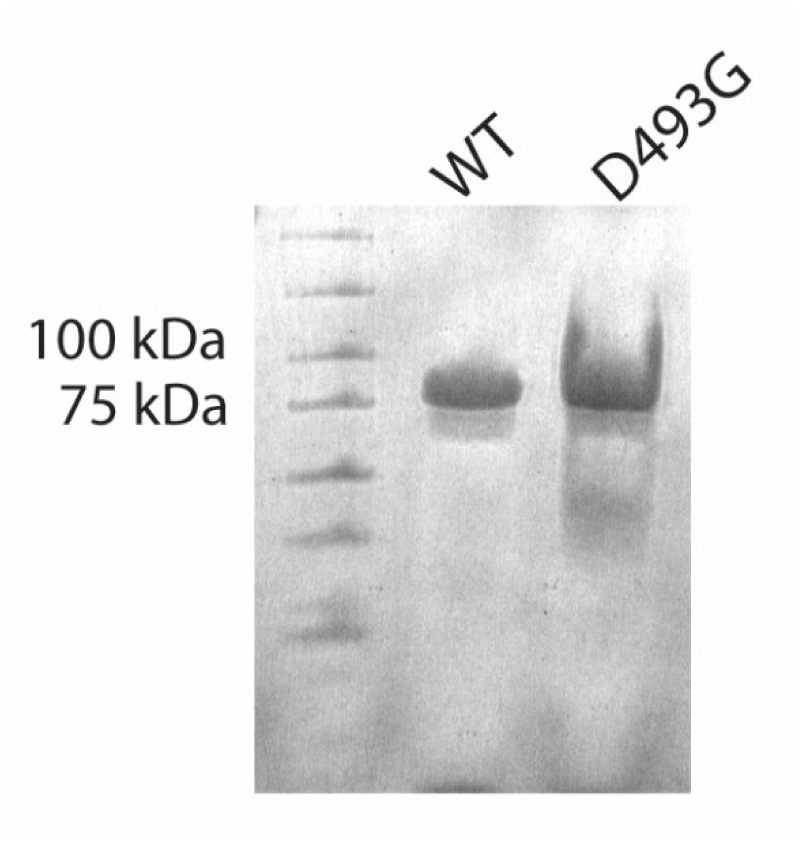
WT and D493G PnpA ran on a 4-20% Mini-PROTEAN® TGX™ Precast Protein Gel (BioRad) and stained with Bio-Safe™ Coomassie Stain (BioRad)

**SI Figure 2.**
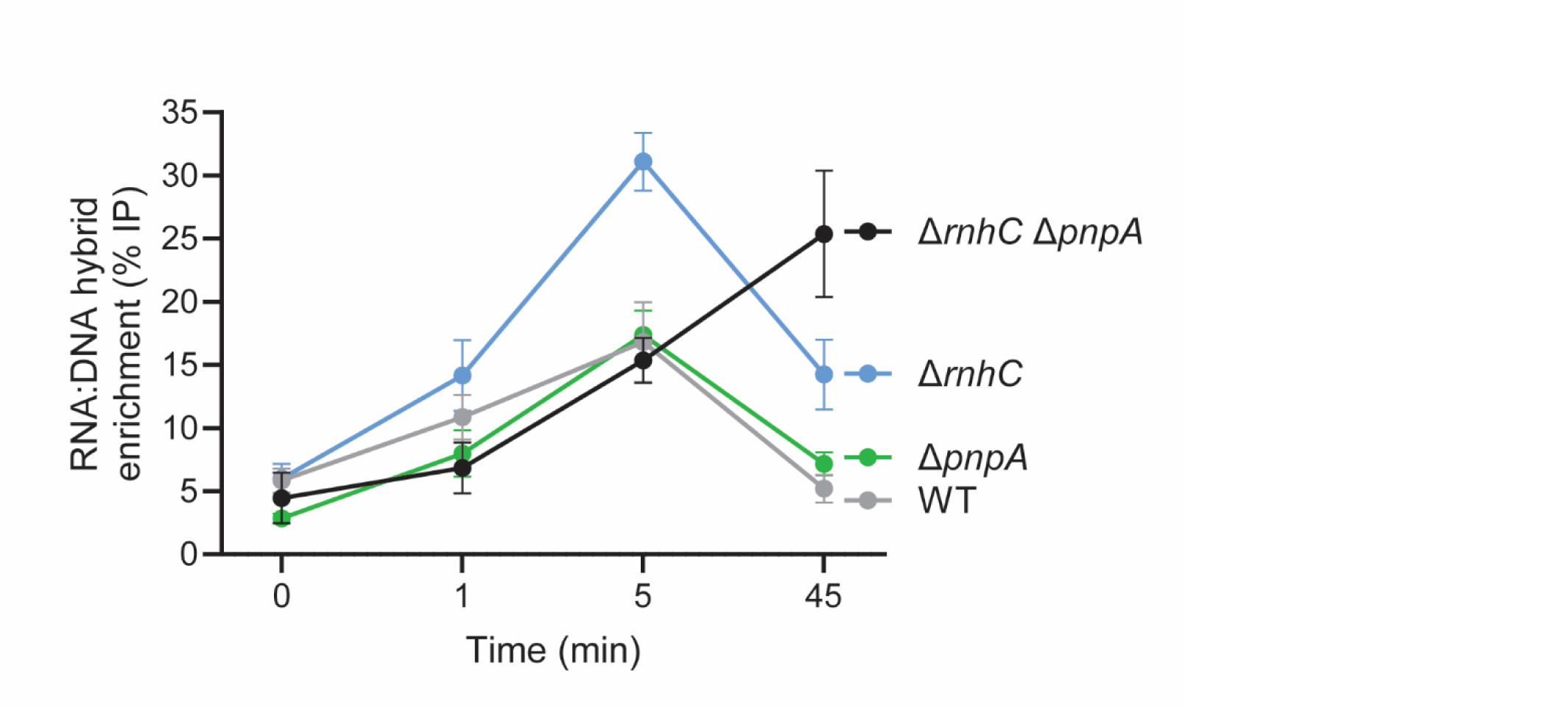
DRIP in the indicated strain represented as % IP before and the indicated time after conflict induction

**SI Figure 3.**
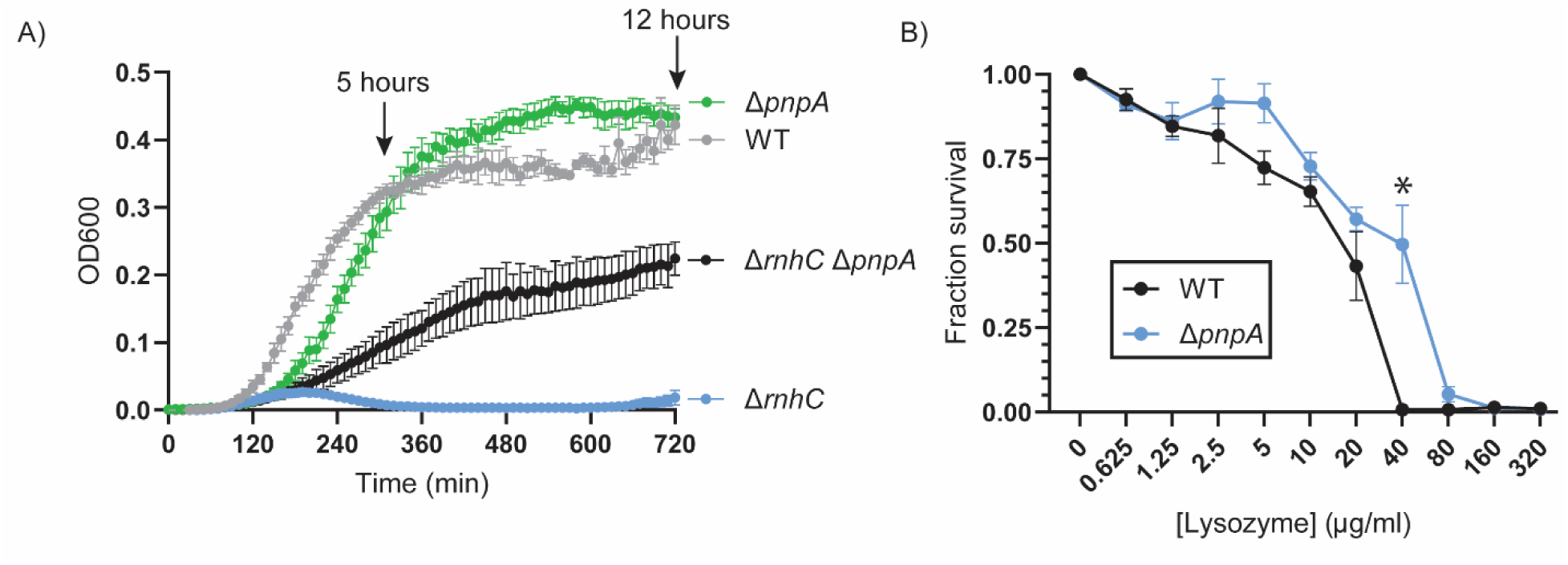
A) OD600 measured every 10 mins for 12 hours in cultures of the indicted strains in the presence of 1 mM IPTG. B) Survival of the indicated strain to the indicated concentration of lysozyme after 16 hours of growth at 37 °C. n=3 biological replicates. Error bars represent the standard error of the mean. Statistical significance was determined by one-way ANOVA *p<.0.05.

### SI Tables

**SI Table 1:** Mutation frequency fold changes and *p*-values.

| Comparison | Fold change | *p*-value (Kruskal–Wallis test) |
| --- | --- | --- |
| WT vs. WT+*rnhC*-OE | 2.0X | 0.25 |
| $\Delta pnpA$ vs. $\Delta pnpA$ +*rnhC*-OE | 2.1X | 0.15 |
| $\Delta rnhC$ $\Delta pnpA$ vs. WT | 6.7X | <0.0001 |
| $\Delta rnhC$ $\Delta pnpA$ vs. $\Delta rnhC$ $\Delta pnpA$ +*rnhC*-OE | 102X | <0.0001 |

**SI Table 2:** Strains used in this study.

| Strain Name | Genotype | Figure |
| --- | --- | --- |
| HM5159 | *amyE::Pspank(hy)-hisC (HO) pnpA-3xmyc* | 1 |
| HM5160 | *amyE::Pspank(hy)-hisC (HO) pnpA-3xmyc rnhC::markerless* | 1 |
| HM452 | *amyE::Pspank(hy)-hisC952-HO* | 2,3,4,S2,S3 |
| HM4257 | *amyE::Pspank(hy)-hisC952-HO rnhC::MLS* | 2,3,S2,S3 |
| HM4275 | *amyE::Pspank(hy)-hisC952-HO pnpA::kan* | 2,3,4,S2,S3 |
| HM4287 | *amyE::Pspank(hy)-hisC952-HO rnhC::MLS pnpA::kan* | 2,3,4,S2,S3 |
| HM2195 | *amyE::Pspank(hy)-luxABCDE* | 3 |
| HM4957 | *amyE::Pspank(hy)-luxABCDE rnhC::markerless* | 3 |
| HM4988 | *amyE::Pspank(hy)-luxABCDE pnpA::kan* | 3 |
| HM4989 | *amyE::Pspank(hy)-luxABCDE rnhC::markerless pnpA::kan* | 3 |
| HM419 | *amyE::Pspank(hy)-hisC952 (HO), hisC952, metB5, leuC427* | 5 |
| HM2482 | *amyE::Pspank(hy)-hisC952 (HO), hisC952, metB5, leuC427 thrC::Pspank(hy)-rnhC* | 5 |
| HM4299 | *amyE::Pspank(hy)-hisC952 (HO), hisC952, metB5, leuC427 pnpA::kan* | 5 |
| HM4305 | *amyE::Pspank(hy)-hisC952 (HO), hisC952, metB5, leuC427 pnpA::kan thrC::Pspank(hy)-rnhC* | 5 |
| HM4303 | *amyE::Pspank(hy)-hisC952 (HO), hisC952, metB5, leuC427 pnpA::kan rnhC::MLS* | 5 |
| HM4309 | *amyE::Pspank(hy)-hisC952 (HO), hisC952, metB5, leuC427 pnpA::kan rnhC::MLS thrC::Pspank(hy)-rnhC* | 5 |
| HM1 | WT | S3 |
| HM4330 | *pnpA::kan* | S3 |

**SI Table 3:**
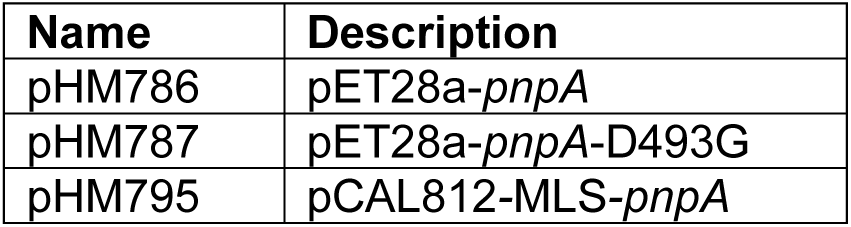
Plasmids generated in this study.

| Name | Description |
| --- | --- |
| pHM786 | pET28a-*pnpA* |
| pHM787 | pET28a-*pnpA*-D493G |
| pHM795 | pCAL812-MLS-*pnpA* |

**SI Table 4:**
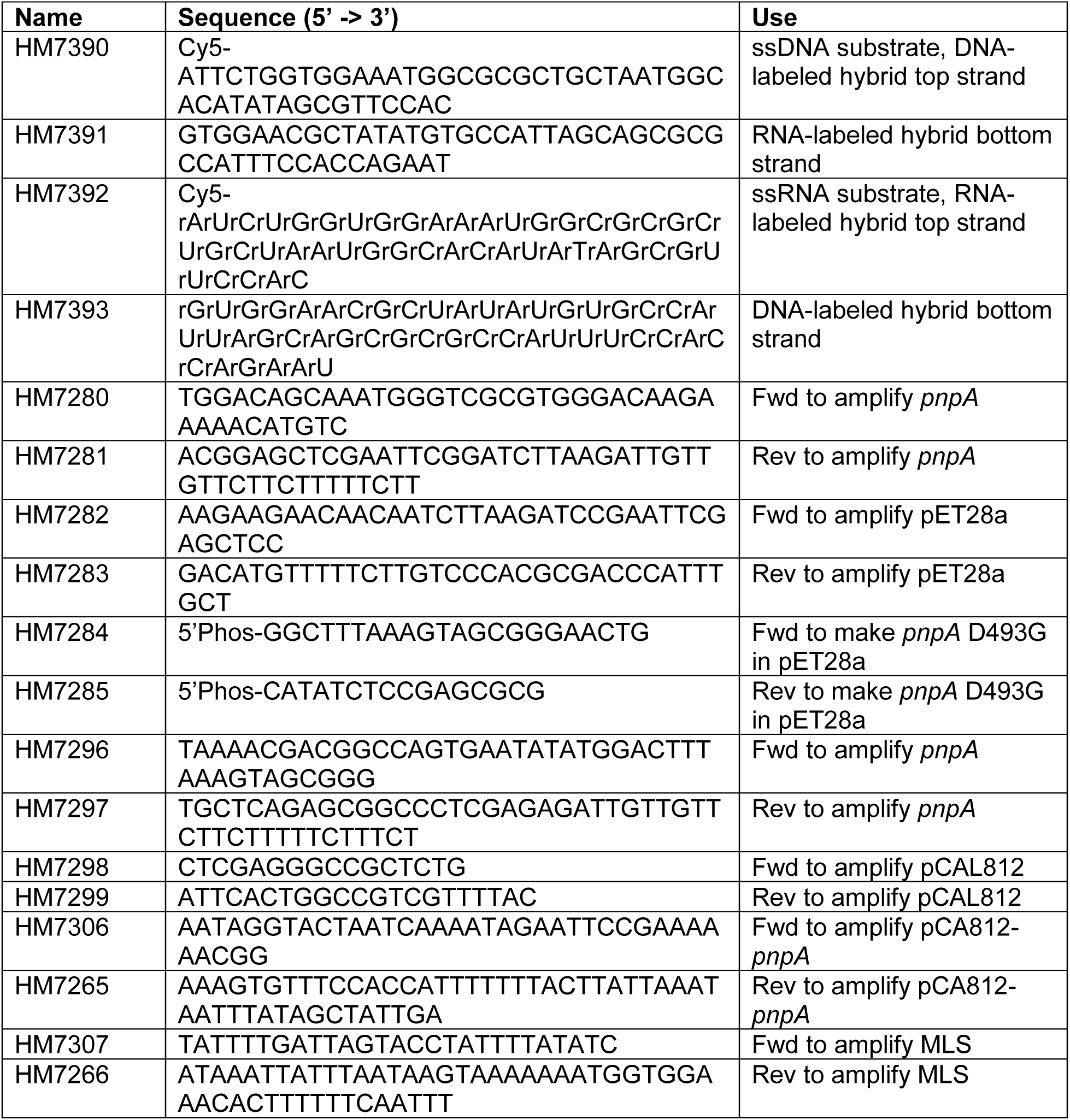
Primers used in this study.

